# A Meta-Analysis of the 16S-rRNA Gut Microbiome Data in Honeybees (*Apis Mellifera*)

**DOI:** 10.1101/2021.12.18.473299

**Authors:** Alexis G. Gkantiragas, Jacopo Gabrielli

## Abstract

1.

Honeybees (*Apis Mellifera)* perform an essential role in the ecosystem and economy through pollination of insect-pollinated plants, but their population is declining. Many causes of honeybees’ decline are likely to be influenced by the microbiome which is thought to play an important role in bees and is particularly susceptible to infection and pesticides. However, there has been no systematic review or meta-analysis on honeybee microbiome data. Therefore, we conducted the first systematic meta-analysis of 16S-rRNA data to address this gap in the literature. Four studies were in a usable format – accounting for 336 honeybee’s worth of data – the largest such dataset to the best of our knowledge. We analysed these datasets in QIIME2 and visualised the results in R-studio. For the first time, we conducted a multi-study evaluation of the core and rare bee microbiome and confirmed previous compositional microbiome data. We established that *Snodgrassella, Lactobacillus, Bifidobacterium, Fructobacillus* and *Saccaribacter* form part of the core microbiome and identify 251 rare bacterial genera. Additional components of the core microbiome were likely obscured by incomplete classification. Future studies should refine and add to our existing dataset to produce a more conclusive and high-resolution portrait of the honeybee microbiome. Furthermore, we emphasise the need for an actively curated dataset and enforcement of data sharing standards.

Figure 1
Graphical abstract.
Made by the author in Biorender.com.

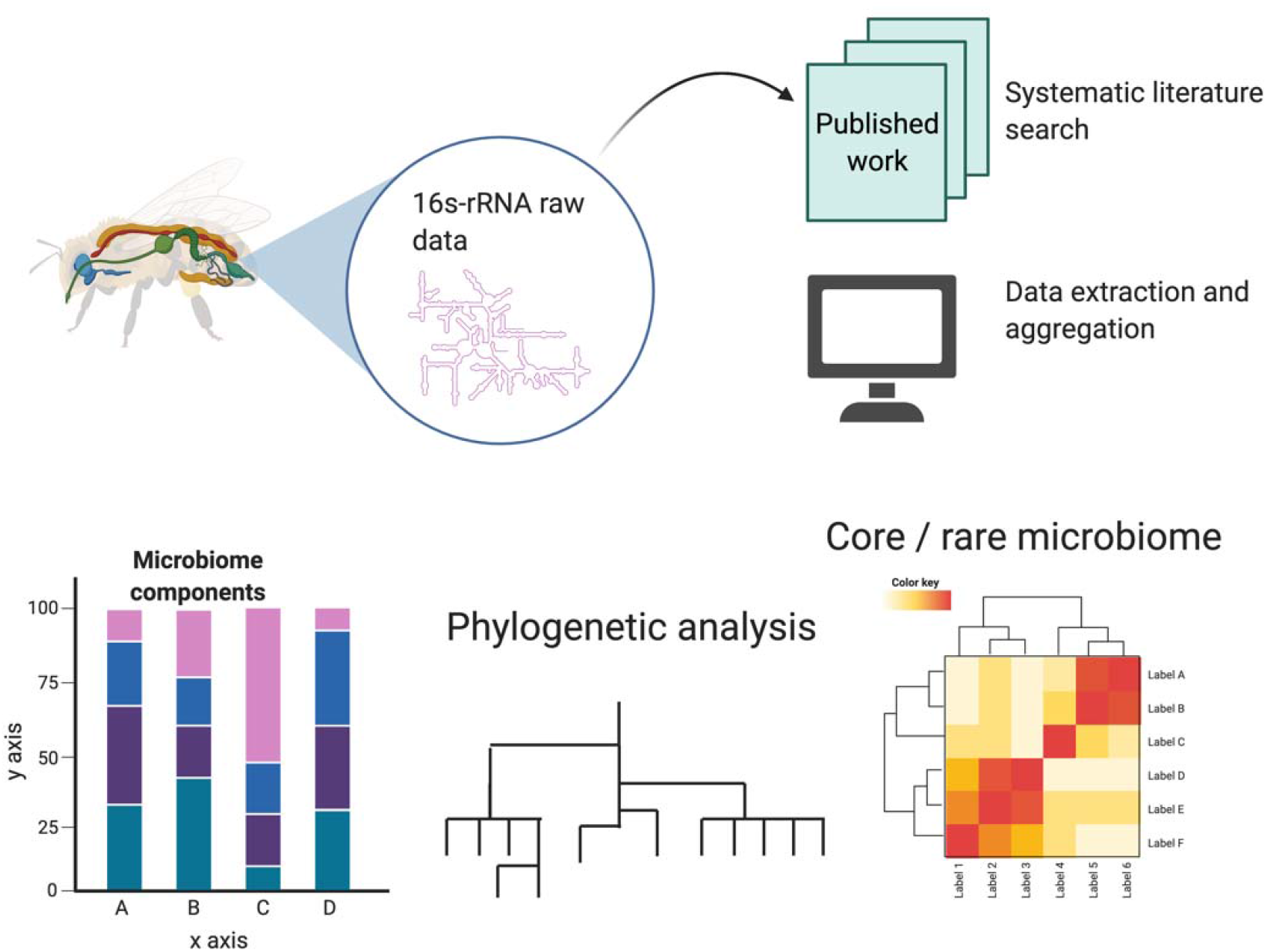

## 2. Introduction

Honeybees (*Apis Mellifera)* play an essential role in the global ecosystem and economy, pollinating most flowering plants and numerous food crops, such as apples and cucumbers^1^. Recent evidence also implies that insect pollinators (honeybees included) perform a role in the pollination of wind-blown pollinated crops, including rice and corn^2^. Thus, the plight of honeybee populations in recent decades^3^ bears grim practical implications.

Aside from loss of flora, the decline in honeybee populations has been attributed to numerous factors — notably, pesticide use and pathogens^4^. These will, to some degree, interact with the microbiome. For instance, the pesticide chlorothalonil was found to lower commensal bacterial sugar metabolism^5^. Similarly, bees live in densely populated colonies with ≥50,000 individuals^6^ and must contest with greater risk from infectious agents. For example, lethal foulbrood infections are caused by the bacterium *Paenibacillus larvae*^7^. In other organisms, symbiotic microbes protect against infection through a combination of outcompeting and direct killing^8^.

A reason honeybees are appealing for microbiome studies is that workers and queens are genetically identical yet display profound morphological and behavioural differences – implying that these are environmentally determined. Individuals become queens when they are fed royal jelly in the early stages of larval development^9^. While the mechanism whereby these developmental changes manifest is still largely unknown, evidence suggests that bacterial colonisation from workers’ glands contributes to the queen-rearing process^10^. Equally, it might also be that the different nutrient composition of royal jelly nurtures different bacterial populations.

While microbiome studies often struggle to demonstrate causative relevance, there is evidence that changes in the microbiome reflect or even dictate phenotypic and behavioural differences in honeybees^11^. Nonetheless, it is worth noting that the hitherto assumed unbiased nature of metagenomics studies have recently been questioned due to substantial bias introduced in DNA extraction and PCR steps^12^.

Bees are generally thought to have a relatively conserved resident population of 8 or so bacterial species^13^. However, these studies principally focused on worker bees, and the microbiome of larvae, drones and queen bees is still largely uncharacterised as it is thought to be more dependent on environmental variation^10,14^. There has not, to our knowledge, been any large-scale validation of these findings to date. The current evidence on the bee microbiome’s role is summarised in an excellent review by Engel *et al.*^15^.

Moreover, most of the individual microbiome members that have been identified have not been linked to specific symbiotic functions within the honey bee gut16. Overall, there is general agreement that the bee microbiome is consistent and stable within workers in a colony which can be attributed both the social nature of bees, the large colony size (up to 50,000 bees per colony), and the use of propolis which works as an antiseptic and reduces bacterial diversity in the hive^16,17^.

Despite the likely importance of the microbiome in bees, relatively few studies have been conducted studying the honeybee microbiome, and there currently exists no meta-analysis or systematic review collating the available bee microbiome data. While the Bee-Biome Consortium has received funding to construct a data portal, this has yet to be completed https://wp.unil.ch/beebiome/. Consequently, this meta-analysis sought to aggregate available datasets on the honeybee microbiome and conduct, for the first time, a multi-study characterisation of the honeybee microbiome. We hope that this aggregated dataset will prove helpful for those conducting future studies.

Although the ‘core’ bee microbiome is referred to in the literature^18^, and can usually be cultured in the lab^13^, this has primarily been summarised in non-systematic reviews and studies. For example, one paper claiming to establish a core bee microbiome contained only 41 honeybees^19^. This leaves such results open to unconscious bias and risks excluding or overrepresenting certain groups of bacteria – particularly as there is no fixed definition of ‘core microbiome’^20^.

We focused upon amplicon sequencing data – a PCR derived approach to perform metagenomic sequencing in which a small sequence or ‘amplicon’ is amplified via PCR and sequenced. Such approaches are essential when studying mixed microbiological samples since most microbes cannot be cultured in laboratory settings^21^. Of equal relevance, amplicon sequencing permits vastly higher throughput experiments.

While more sophisticated methods such as whole-genome next-generation sequencing (NGS) have become more widespread in microbiome studies, including in bees^22^, the examination of such methods was beyond this study’s scope. Though the cost is ever decreasing, these methods are more expensive and demand a more complex workflow. Nonetheless, future studies may wish to consider incorporating such data into a metaanalysis.

## 3. Methods

### 3.1. Systematic Search

Firstly, we performed a systematic review of the literature on PubMed using the search terms: *(microbiome AND “16S rRNA” AND “Apis mellifera”)*

This search was performed on 22/10/2020 yielding 26 results. Next, searches for data availability were performed independently by two researchers. A total of 14 studies were excluded on the basis of lack of available data or data available in an unusable form (e.g. in PDFs). This left a total of 12 studies to be used in the meta-analysis. The annotated paper database can be found in our Github repository (see data availability statement).

Of the 12 studies where data was accessible, most were either multiplexed and with no barcode file to allow demultiplexing, had imprecise structure and/or metadata or had considerably low-quality sequences, making them hard to extract and analyse. Finally, only four studies had a clear data structure and were fully accessible and reproducible on QIIME2. Accordingly, these studies were used subsequently and are detailed below.

We did not intentionally include studies on bumblebees or other related species as, while valuable, they were beyond the scope of this study. Additionally, papers that did not have experimental data available publicly were excluded due to time constraints on this study. We also excluded studies investigating *nosema* infestation as we reasoned this would likely perturb normal microbiome function. Nonetheless, these would not be picked up since bee parasites such as *nosema* are eukaryotes, and *nosema* infestation may be asymptomatic^23^. Studies investigating the specific effect of *nosema* infestation on microbiome function and corresponding meta-analysis would be valuable additions to the literature.

We focused upon 16S-rRNA data since this is a well-established bacterial marker used routinely in amplicon metagenomics sequencing. We used the GreenGene database for the identification of 16S-rRNA https://greengenes.secondgenome.com/. Pipeline Files in .sra format were downloaded from SRA, where they were converted to FASTA format, readable by QIIME2 and aggregated. Various data types were obtained; thus, adjustments and labelling had to be carried out.

#### 3.1.1. A – Saelao *et al.*, 2020^17^

The authors studied 62 bees from 32 colonies. They used an equal number of bees from propolis-rich and propolis-poor colonies and concluded that the former was associated with a ‘stabilised’ microbiome. However, they did not link these to any colony outcomes and neglected to describe how propolis richness was determined.

#### 3.1.2. B – Horton *et al.*, 2015^24^

This paper claimed to define and confirm earlier experimental findings of the ‘core’ bee microbiome. Unfortunately, they did not clarify the number of samples used in their experiments – only 35 were present in the dataset.

#### 3.1.3. C – Erban *et al.*, 2017^25^

This study used 49 bees covering 27 colonies. Of note, some of the colonies in this sample were affected by foulbrood, likely accounting for this genera’s presence within the rare bee microbiome, as examined later.

#### 3.1.4. D – Powell *et al.*, 2016^26^

This study contained 221 samples from an unspecified number of colonies. Unfortunately, we failed to exclude some bumblebee samples from this group, possibly accounting for the large number of outliers in D, as discussed later.

### 3.2. Diversity Metrics

Measures of alpha diversity were performed to ascertain each study’s microbial diversity and the variation in diversity. Shannon diversity (Equation 1)(first described in 1948^27^) was used as a non-phylogenetic measure of alpha diversity, while faiths-PD (Equation 2)(first described 1992^28^) was employed as a phylogenetic measure of alpha diversity. Statistical significance was determined using Turkey’s test^29^.

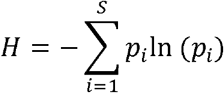

*Equation 1 - Shannon diversity index. Wherein*:

*H = Shannon diversity index*
*S = Number of species*
*P_i_ = Proportion of abundance represented by i^th^ species*

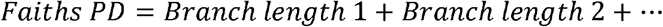

*Equation 2 - Faiths PD. Calculated via the sum of branch lengths from a phylogenetic tree.*

Beta diversity was assessed in a PCoA plot via a Jaccard index (first developed 1912^30^) as shown in Equation 3 and Bray-Curtis dissimilarity matrix (first developed 1957^31^), shown in Equation 4. PERMANOVA^32^ was used for comparisons of beta diversity.

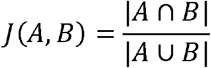

*Equation 3 - Jaccard similarity matrix.*

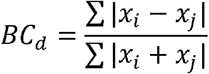

*Equation 4 - Bray-Curtis dissimilarity matrix. Wherein*:

*BC_d_=Bray-curtis dissimilarity value*
*x_i_=abundance of species in sample i*
*x_j_=abundance of species in sample j*

### 3.3. Data Visualisation and Analysis

Analysis was performed in QIIME2, and visualisation of data was performed in RStudio due to its superior processing times compared to similar pipelines^33^. A QIME2 pipeline (Figure 2) was utilised as described previously^34^. We utilised both DADA2^35^ and the QIME2 deblur function for the denoising step due to difficulties in aggregating multiple datasets. We used 97% OTU clustering.

**Figure 2.**
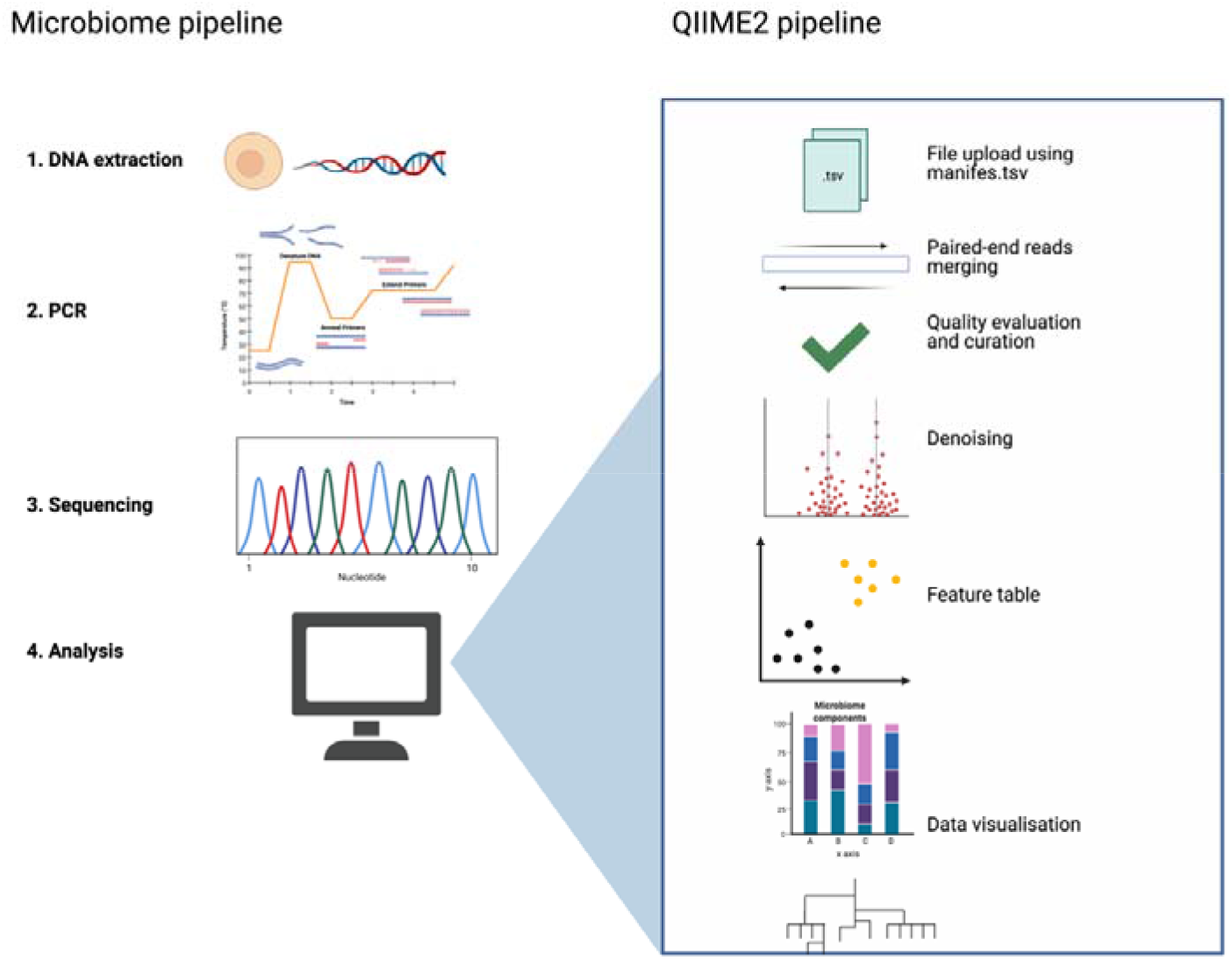
Overall microbiome study- and QIME2 pipeline. We conducted only the microbiome pipeline’s final step – that is, analysis due to lack of access to a laboratory space. Made by the author in Biorender.com.

We used several libraries to handle and visualise the data. These are described in our Github repository https://github.com/JacopoGabrielli/Bee_Microbiome?fbclid=IwAR0AvpfowNnWHNDlFXM5z24a6ws9f5zKmF9o1n8DRXWH9SLqJ-YPHQo4qVg.

## 4. Results

### 4.1. Diversity Measures Imply a Consistent Microbiome

We began by measuring alpha diversities of each study to confirm previous findings that the honeybee microbiome is relatively consistent. Samples were rarefied to 1300 reads to provide sufficient coverage.

All studies except A and B had statistically significant differences in non-phylogenetic alpha diversity as measured by Shannon diversity (Figure 3A). Even A/B have such large error bars that the result might be obscured by the sizeable signal-to-noise ratio. This suggests that the microbial diversity varied across studies. Likewise, there appears to be a great deal of intra-study variation between bees, with data points in group A showing almost 10-fold differences in diversity. This implies that individual bees possess vastly dissimilar microbial diversity. Overall, it is worth noting that these are somewhat low alpha diversity values.

**Figure 3.**
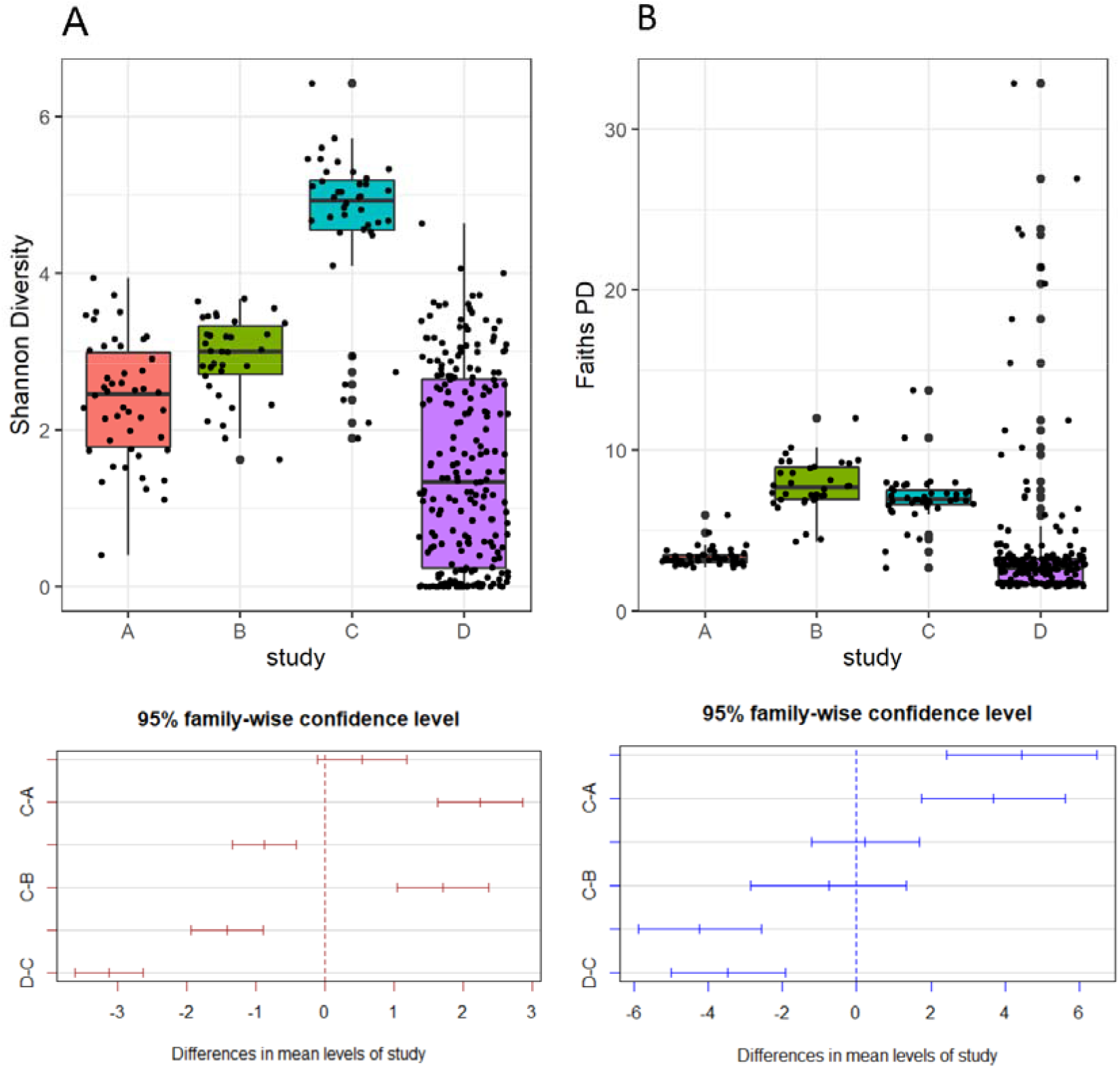
Differences in diversity between bee microbiome studies. A) Shannon diversity and comparison between studies (shown below). B) Faiths-PD values for each study and comparisons between them. Error bars show extreme upper and lower bounds while boxes represent upper and lower quartiles with the centre line’s median. Each data point shows a single sample (Bee). 95% family wide confidence levels were calculated using Turkey’s test. For A-B the box plot shows study letter/code on the x-axis and the diversity index on the y-axis.

Once adjusting for phylogenicity using Faiths-PD, this picture changes (Figure 3B). Most noticeably, the intra-study variation appears significantly diminished, with comparatively tiny error bars for each study. However, this might be a result of the altered scale. While D contains many outliers at approximately 10-fold the mean, this is presumably a product of the large sample size D used. It is interesting that these outliers only become apparent once phylogeneticity is accounted for. These results suggest that their variation in the bee microbiome primarily occur between related microbial species.

Limited information can be divined from a PCoA plot of the studies. While C and D showed overlapped, all other studies showed no similarity to each other. This applies both when using a Jaccard- (Figure 4A) and Bray-Curtis similarity matrix (Figure 4B).

**Figure 4.**
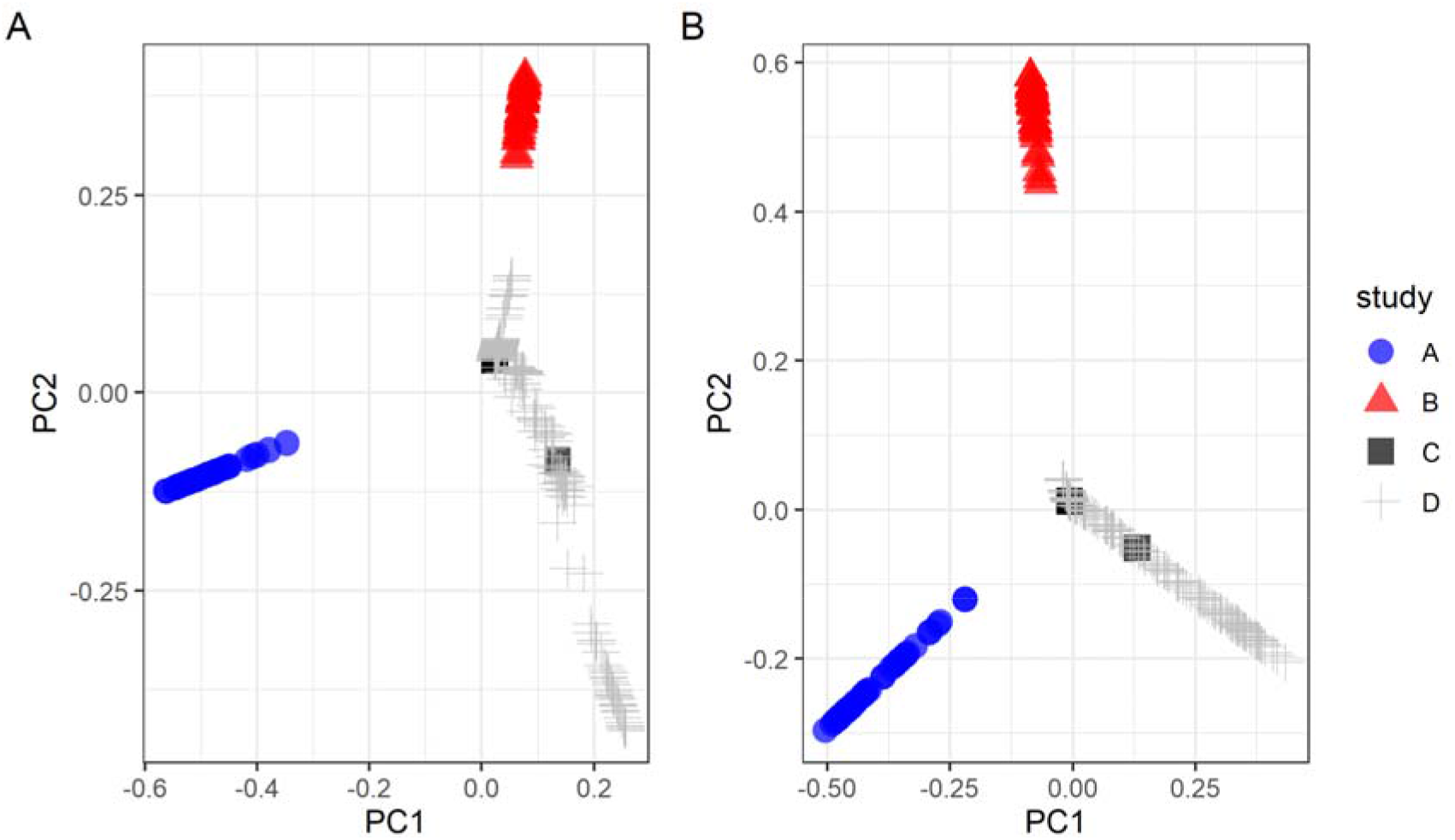
PCOA plot. using A) Jaccard similarity matrix. B) Bray-Curtis dissimilarity matrix. For A) and B), the x-axis represents PC1 which accounts for the greatest variance in the data while the y-axis represents PC2 which accounts for the second greatest variance in the data. Each data point represents an individual sample within its respective study.

### 4.2. Taxonomic Classification and Composition

Following phylogenetic analysis, the only class identified was alphaproteobacteria, though many samples lacked an applicable class (Figure 5A). Oddly, these are spread across various clades instead of grouped under one. It is also worth noting that there are not many clades in the phylogenetic tree. Likewise, branch length is relatively short, and there is minimal branching (Figure 5A). This would speak to the consistency of the honeybee microbiome.

**Figure 5.**
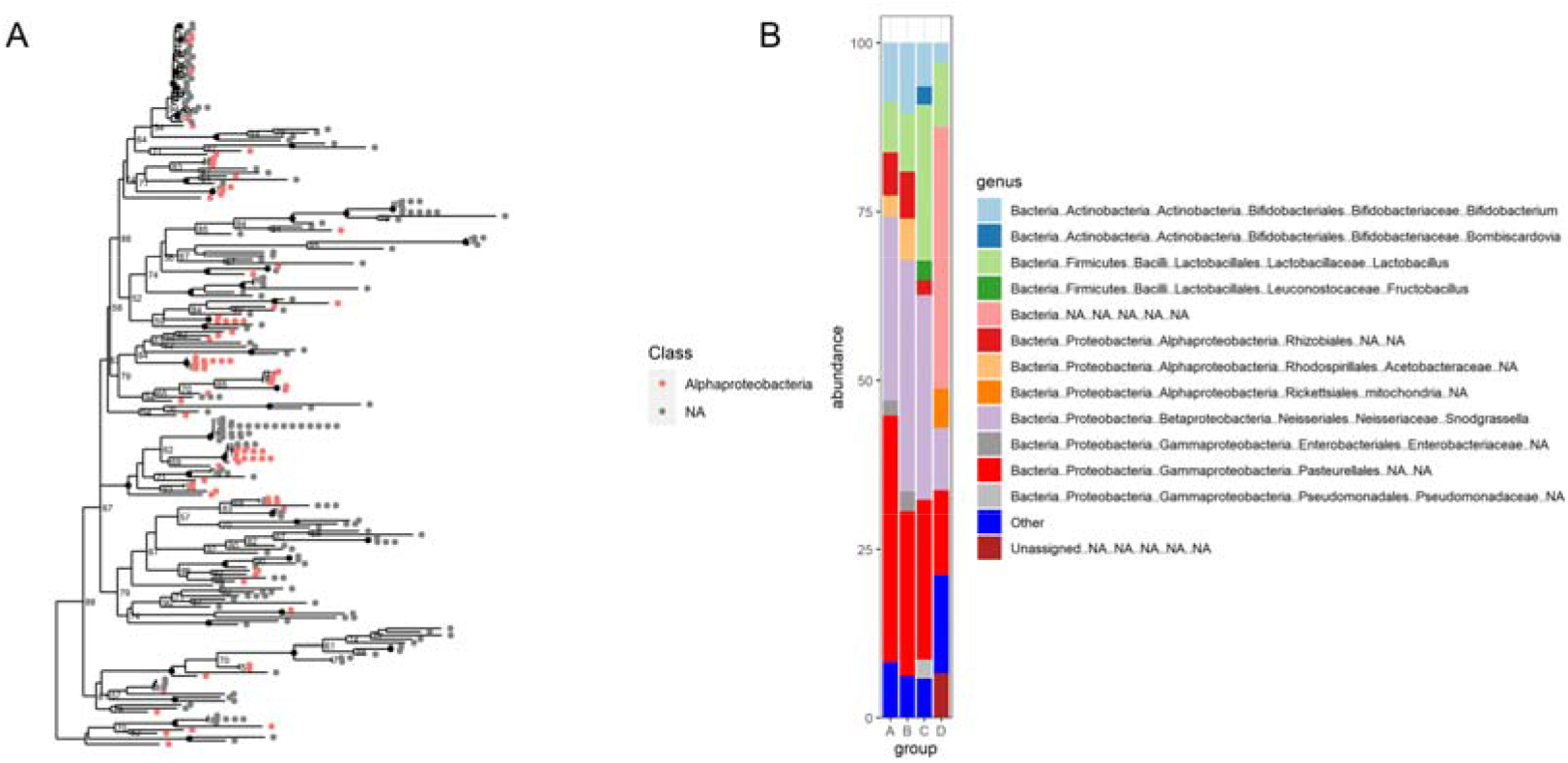
Composition and phylogeny of the bee microbiome. A) phylogenetic analysis. Dots represent individual taxa. The numbers are branch support values; they are a percentage reliability measure of separation from surrounding clades/branches B) Genus composition of studies A)-D). In B) the x-axis represents the group/study letter while the y-axis shows the abundance percentage.

Four genera dominate across all studies and account for most of the abundance in each (Figure 5B). The abundance of these groups is relatively constant across every study. These comprise 1) *Bifidobacterium,* 2) *Lactobacillus,* 3) *Snodgrassela,* and finally, 4) (unknown genus) the order *Pasteurellales.*

*Bifidobacteria* are gram-positive species found in the human gut and frequently used as probiotics^36^. These comprised around 10% of samples in A-D. These have also been reported in various bee digestive systems, including bumblebees^37^, honey bees^38^ and carpenter bees^39^. They are broadly thought to have a beneficial effect on bee health, as they do in humans. Indeed, dietary supplementation of *Bifidobacteria* was protective against *nosema* infection in honeybees^40^.

Similarly, *Lactobacillus* are a gram-positive genus used as a probiotic and found in the human gut^41^. Indeed, in honeybees, these buffer against antibiotic-induced dysregulation^42^. Moreover, their supplementation to honeybee colonies (in *Lactobacillus johnsonii)* improved egg laying capacity^43^.

*Snodgrassella* bacteria were originally isolated in honeybees, and their function remains unknown^44^. Finally, the order *Pasteurellales* are most frequently found on mucosal surfaces and are gram-negative bacteria^45,46^. They are believed to have a broadly symbiotic function^47^.

Overall, we can see that the bacterial bee microbiome appears to be dominated by commensal, gram-positive species whose supplementation promotes health. Since honeybees exist in huge colonies, it is unsurprising that they would have conserved commensal microbiota, which protects against the higher risk of infection. It also may be a way to help ensure stability and consistency across individuals. While the main genera are all shared across studies, the incomplete classification of many taxonomic groups makes the results challenging to interpret.

The presence of *Bombiscardovia* in C is likely a result of contamination with bumblebees since this genus is exclusively found in bumblebees^48^. Indeed, after revisiting the dataset, it appears that we failed to exclude bumblebee samples. It is also worth noting that many of D’s species (which were also the largest group) are either unidentified or unassigned.

Other studies reported *Bartonella*^49^ and *Gillamella* genera which we did not observe. However, this is doubtless due to poor sequencing information as *Bartonella* can likely be accounted for by the “Rhizobiales...NA...NA” (for it falls within this order). Interestingly, in the case of *Gillamella* – past studies utilising QIIME had noted that it mislabelled *Gillamella* and *Frischella* as *“Pasteurellales”*^50^. This is somewhat unusual given that *Gillamella* does not belong to the order *Pasteurellales*.

### 4.3. The Core and Rare Microbiome

We attempted to characterise the core (Figure 6A) and rare (Figure 6B) honeybee microbiome. Most obvious is that ‘unknown’ comprise the most common group in the data. This implies that the bee microbiome is poorly characterised. The difference between Figure 6 and Figure 5 can largely be accounted for by the large number of unknowns (labelled NA in the previous graph) and the data’s heterogeneity. Furthermore, group D is far more heavily weighted here owing to the larger sample size. Nevertheless, the same genera are represented here, such as *Snodgrassela, Lactobacillus* and *Bifidobacterium.*

**Figure 6.**
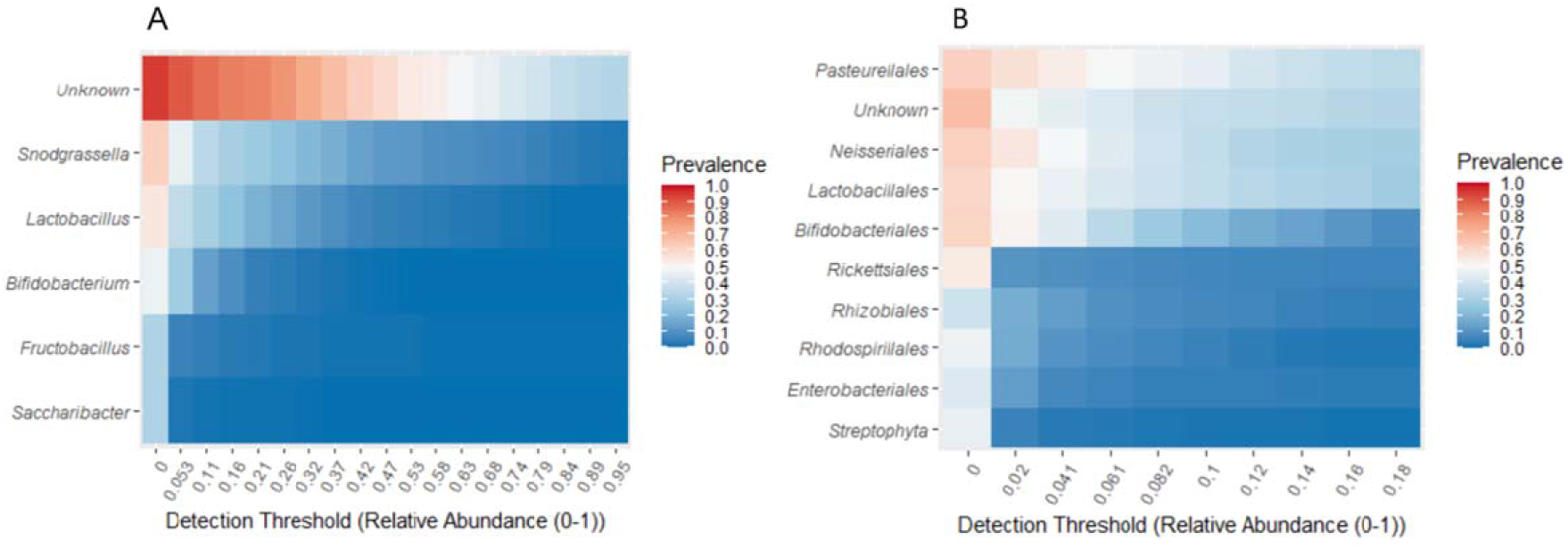
Heatmaps of the honeybee microbiome. This represents a weighted and aggregated form of the four studies. A) The core bee microbiome. These were sorted in order of prevalence. B) The core microbiome at order level. The y-axis indicates the genus (A) and order (B) while the x-axis denotes the detection threshold in A-B.

Interestingly, *Fructobacilla* and *Saccharibacter* are both recently discovered gramnegative genera^51^. *Fructobacilla* are, as the name suggests, a form of lactic acid bacteria which thrive in fructose-rich environments and have been well described in bee guts^52^. *Saccharibacter* are a genus with presently only one species (*Saccharibacter floricol*) which has been described in honeybees^19^ and was first isolated from flower pollen^53^ – from which one might speculate the guts were colonised. They both play a role in sugar metabolism – suggesting a probable commensal function as microbes often aid in digestion^54^.

In considering the core bee microbiome orders (Figure 6B), we see that various unusual bacteria are present, including *Rhizobium,* most well known as a nitrogen-fixing bacteria found in the root nodules of legume plants^55^. These are likely *Gillamella* which belong to the order *Rhizobium*.

A total of 251 rare genera were identified. These were filtered by setting a maximum prevalence of 3%. Interestingly, this included 8 genera of archaea which were primarily involved in methane processing. This may have been due to improper storage or preservation of the bee samples, which may have led to decay in one or two instances. This occurs rapidly for a small organism like a honeybee. Nonetheless, archaea have been reported (albeit at very low abundances) in Africanized honeybees^56^.

Finally, two *Paenibacillus* genera were detected in the rare microbiome, a known honeybee pathogen (American foulbrood)^7^. This is likely the result of foulbrood samples within C – though recent studies reporting asymptomatic foulbrood^57^, *nosema*^23^, *crithidia*^58^ and chronic paralysis virus^58^ presence do raise the question of whether *Paenibacillus* might have been present in other studies. A more complete bee microbiome dataset might clarify this issue – particularly given that this would have significant impacts for both wild and captive bee management.

Differential abundances for classes of bacteria were conducted to evaluate these groups’ abundance between studies. Interestingly, study A showed the greatest variation in samples (bees) in abundance (Figure 7). Proteobacteria in study A had outliers approximately 4-fold higher than those of any other groups. These are gram-negative bacteria which include Snodgrassela and Gilliamella. We see that there is a greater median abundance in A and B. Additionally, A and B als show far greater variation. For visualisation purposes, a single outlier in B was excluded.

**Figure 7.**
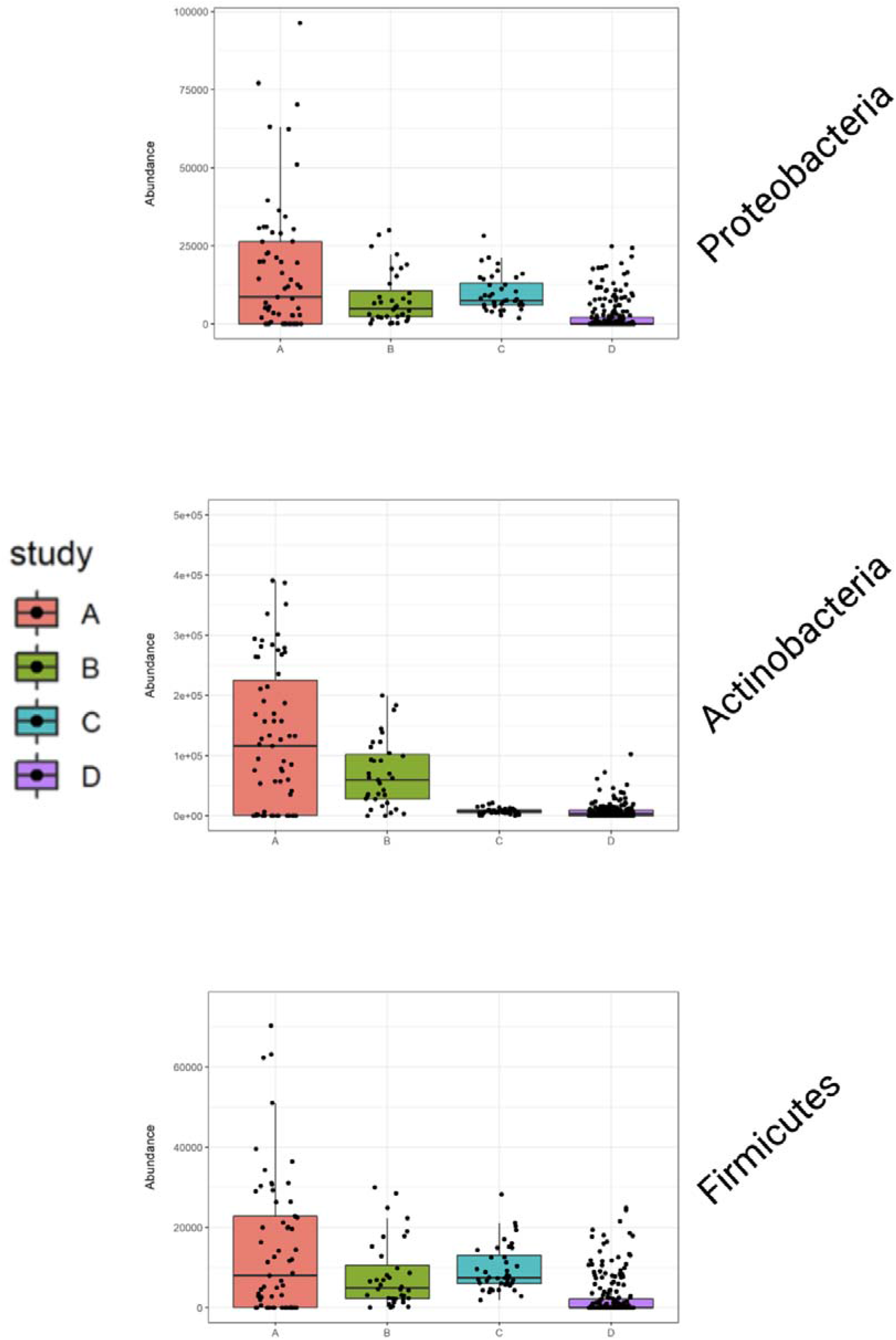
Differential abundance for the classes proteobacteria, actinobacteria and firmicutes. Error bars show extreme upper and lower bounds; boxes show upper and lower quartiles with the median represented by the centre line. Individual data points represent individual bees. The y-axis denotes abundance (though scales vary) and the x-axis shows the study letter/code.

Again, *Actinobacteria* were more abundant in study A (Figure 7). These are a grampositive phylum of bacteria frequently found in bee guts and have even been found to produce compounds that inhibited *Paenibacillus larvae*^59^. Although this study was conducted in *Melipona scutellaris* (Stingless bees), the results are likely translatable to honeybees. These correspond to orders such as *Streptomyces* noted in the core bee microbiome (Figure 5B). Oddly, these are generally thought to have higher abundance than we obtained^60^.

*Firmicutes’* differential abundances appear to be relatively homogenous across all samples except for D, which has a very low median (Figure 7). This is quite strange and might be due to impurities or sequencing issues since this group had a large amount of NA data. There was also greater variation in abundance between samples in A with several outliers showing around 10-fold the abundance versus the median. These correspond to genera such as *Lactobacillus*.

## 5. Discussion

### 5.1. Agreement with Prior Studies

Overall, our results confirmed previous studies’ findings, confirming the ‘core’ bee microbiome in the first meta-analysis in this area. However, we also highlighted the need for further classification of the bee microbiome and failed to replicate some of previous studies’ findings, such as the prevalence of *Gillamella.*

### 5.2. Limitations

Our investigations had several constraints. Unfortunately, we were unable to access data from most available studies. Similarly, data that required laborious mining techniques could not be used due to time constraints. Searches on Web of Science and other databases could have been searched but were not due to material and time constraints on the study. The authors will conduct these at a later date.

Due to investigating only 16S-rRNA, we were unable to study viruses, prions or fungi – which nonetheless play a role in bee health^61,62^. However, these should be studied separately since these would require sequencing different regions and consequently, datasets which contained all types of microbes would likely be too small. Similarly, we did not look into specific subset populations like the bees’ surface and the specific composition of each body part e.g. malpigian tubules vs hindgut and honey crop (Figure 8). This is because many studies use whole bee samples – thereby prohibiting organ-based differentiation. While subsequent studies may wish to take such a differential approach, this would require skilled and time-consuming dissections.

**Figure 8.**
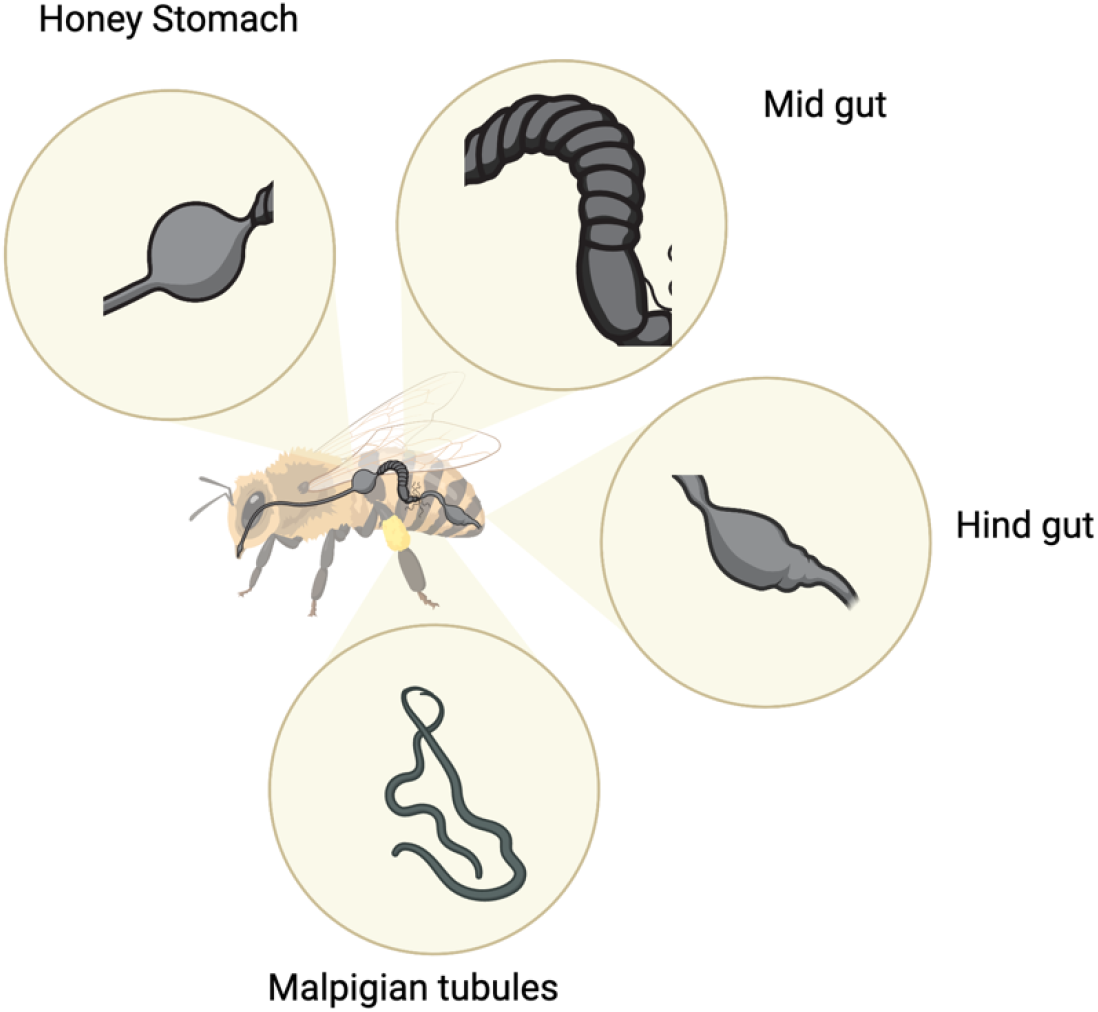
Honeybee worker digestive anatomy. Shown enlarged are the major components of the bee digestive system, which would be well suited to closer study. Made by the author in Biorender.com.

Many different measures of microbial diversity could have been used. In particular, studying phylogenetic beta diversity, such as via UniFrac analysis^63^, would be a worthwhile addition since we only used non-phylogenetic measures. This might increase clustering between groups, given that variation in alpha diversity appeared reduced upon using Faith’s-PD.

Moreover, due to the nature of meta-analyses, we have no way of knowing whether contamination in the samples may have occurred. We must simply hope that these do not directionally bias the results of our study.

Finally, since this is an active research area, several studies (such as Ptaszyńska *et al.,* 2021^64^) were added throughout our analysis that would have fulfilled inclusion criteria. This underlines the importance of a curated and active database as proposed by Engel *et al.* We intend to include all studies published since the initial search in a later iteration of this study.

### 5.3. Future Directions

Subsequent studies should consider contacting authors of papers with registered but uninterpretable datasets, checking for papers published since the initial literature search and finally conducting similar meta-analyses for different bee types (e.g. bumblebees). This approach should be easily translatable to similar insect types. In the future, we aim to improve our dataset by adding recently published papers and constructing a web app to make the data more accessible.

Given the ever-increasing affordability of genetic sequencing, a well-characterised bee microbiome may have eventual commercial relevance in smart bee monitoring. This could provide an additional dimension to be considered alongside hive temperature, humidity and honey production^65^.

Similarly, while various isolated incidents of asymptomatic infections (described earlier) have been reported, such findings can only truly be evaluated by large scale meta-analysis to ensure that the results were not simply anomalous. If chronic infections are prevalent in bees, this could shift emphasis of bee management towards general maintenance of hive health as opposed to control of infection sources. Moreover, if, as one study of over 1000 bees suggested^58^ chronic asymptomatic infections are quite literally ubiquitous in bees, it would be more proper to consider them a resident component of the bee microbiome and regard them as opportunistic pathogens. Thus, future studies should avoid excluding infected bees.

While incredibly ambitious, one might consider constructing a time-course-based metaanalysis, matching bees by age, thus assembling a picture of the bee microbiome over the life course. Although huge datasets would be required for such subgrouping, it might provide enhanced resolution of the bee microbiome and provide novel insights into bee physiology.

Finally, a potentially exciting area for study would be creating a separate ‘wild’ honeybee core dataset. Wild bees have different foraging habits^66^ and have overall smaller, less densely packed colonies — particularly as captive bees are kept in apiaries with many hives close together.

## 6. Conclusions

While significant evidence supports the importance of the microbiome in bees, its true relevance cannot be determined until the standard bee microbiome has been elucidated. Once established, deviations from this core microbiome, and indeed all insights in bees’ microbiomes will gain greater relevance. Unfortunately, there is presently no standard core or rare microbiome in the honeybee. We have established, for the first time, a core honeybee microbiome though further characterisation is needed. To facilitate future research in the field, we have made this publicly accessible to the community as a reference. Our metaanalysis demonstrates that there is a need for standardisation and enforcement of data sharing policies. Specialist journals in the field would be well advised to require accessible data sharing.

## 7. Data Availability and Conflict of Interest Statement

All data is freely and publicly available on Github (https://github.com/JacopoGabrielli/Bee_Microbiome?fbclid=IwAR0AvpfowNnWHNDlFXM5z24a6ws9f5zKmF9o1n8DRXWH9SLqJ-YPHQo4qVg). Both authors read and approved the final manuscript.

Alexis Gkantiragas is a voting member of the BeeXML standardisation Apimondia working group, though does not receive financial compensation for this involvement.

## 8. Acknowledgments

We would like to thank Dr Toryn Poolman his assistance with this study. Furthermore, we would like to thank the colleagues who helped review this manuscript.

